# Pathogenicity of novel *Monosporascus* species in accessions of four melon botanical groups

**DOI:** 10.1101/2022.07.28.501868

**Authors:** Francisco Cleilson Lopes Costa, Andréia Mitsa Paiva Negreiros, Rui Sales, Glauber Henrique de Sousa Nunes

## Abstract

The Monosporascus root rot and vine decline (MRRVD) can be caused by the fungal species *Monosporascus cannonballus* and *M. eutypoides*. The *Monosporascus* species recently described in Brazil (*M. brasiliensis, M. caatinguensis, M. mossoroensis, M. nordestinus* and *M. semiaridus*) are potentially causal agents of the MRRVD. This work was made to evaluate them comparing to *M. cannonballus* pathogenicity, and to evaluate melon accessions reaction in four botanical groups of melon. MRRVD was evaluated by the severity of the damages in the roots and by the root dry matter reduction index (RI_DM_). On average, the studied species caused damage to melon accessions. After all, only *M. brasiliensis, M. caatinguensis* and *M. nordestinus* were virulent according to the accessions evaluated (A-16, C-32 syn. Pat 81, ‘Goldex’ and ‘HBJ’), i.e., accession-species interaction occurred, and among them, *M. caatinguensis* was the most virulent. The accession A-16 (*acidulus* group) showed higher resistance (<10 % of root dry mass loss) to *M. cannonballus, M. caatinguensis* and *M. nordestinus*. The accession C-32 was susceptible to *M. caatinguensis* and moderately resistant to the others. The accession A-16 was the most promising one and can be used as a donor of resistance alleles or as a rootstock.

## Introduction

The MRRVD is a cucurbits syndrome caused by many soil-borne fungal species. To date, in the *Monosporascus* genus, only *M. cannonballus* Pollack & Uecker (Pollack and Uecker 1974) and *M. eutypoides* (Petrak) von Arx were reported as an MRRVD causal agents. The syndrome shows no visible symptoms up to the fruit ripening, that comes two weeks about the harvest, whereas the vines show a yellowing as reflex of resistance to the hydraulic conductance throughout the xylem vessels caused by the root rot. A defoliation progressively occurs after the vines yellowing up to a partial or complete decline of the vines (Cluck et al. 2009; Picó et al. 2008). MRRVD have been causing many losses worldwide, mainly in melon (*Cucumis melo* L.) and watermelon (*Citrullus lanatus* [Thunb.] Matsum. & Nakai) (Iglesias et al. 2000a; Sales Júnior et al. 2012; Al-Mawaali et al. 2013; Yan et al. 2016; Markakis et al. 2018; Negreiros et al. 2019).

Among the *Monosporascus* genus, only *M. cannonballus* and *M. eutypoides* (Petrak and Ahmad 1954; Ben Salem et al. 2013) were reported as MRRVD causal agents (Castro et al. 2020). From these to now, Negreiros et al. (2019) reported five novel *Monosporascus* species associated to two weeds (*Boerhavia diffusa* L. and *Trianthema portulacastrum* Linn) in northeastern Brazil. These species are *M. brasiliensis, M. caatinguensis, M. mossoroensis, M. nordestinus* and *M. semiaridus*, of which A. Negreiros, M. León, J. Armengol and R. Sales Júnior are the authorities of all them. We hypothesize these species as potential causal agents of MRRVD to melon, anyhow to our knowledge there is no works proving this hypothesis to date.

Several works reported melon genotypes with moderate to high resistance to *M. cannonballus* (Salari et al. 2012; Park et al. 2013; Sales Júnior et al. 2019). Some of them surveyed potential genotypes for melon breeding programs, for selection of melon genotypes (Park et al. 2013) and pumpkin genotypes for root stocks (Ben Salem et al. 2015a) and selection of resistant melon cultivars (Salari et al. 2012; Sales Júnior et al. 2019). These researches demonstrate the MRRVD genetic control is one of the best alternative solutions for controlling *Monosporascus* species. The sources of resistance of *C. melo* to MRRVD shall have high root vigor and high heterosis of root traits (Cohen et al. 2012b).

Thus, to our knowledge sources of resistance to *Monosporascus* species described in Brazil are unknown and must be investigated to support melon breeding programs for affording the production of MRRVD resistant cultivars. We surveyed four melon genotypes for resistance to six *Monosporascus* species. The objective of this work was to evaluate the novel *Monosporascus* species pathogenicity comparing them to *M. cannonballus* and to identify melon accessions resistant to MRRVD.

## Material and Methods

### Plant Material

The melon accessions are two commercial cultivars (‘HBJ’ - *Hales Best Jumbo*®, ‘Goldex’®) and two genotypes A-16 and C-32 (syn. Pat 81) that are part of the cucurbits germplasm bank of Universidade Federal Rural do Semi-Árido (UFERSA-Brazil). The ‘HBJ’ accession belongs to the *cantaloupensis* botanical group, the ‘Goldex’ accession belongs to the *inodorus* group, the accession A-16 belongs to the *acidulous* group and C-32 belongs to the *conomon* group.

### *Monosporascus* isolates

The fungal isolates utilized in this work are stored in the fungal collection of Phytopathogenic Fungi “Prof. Maria Menezes” (CMM) at the *Universidade Federal Rural de Pernambuco* (*Recife, Pernambuco*, Brazil) with the following codes *M. brasiliensis* (CMM 4839), *M. caatinguensis* (CMM 4833), *M. cannonballus* (CMM 2386), *M. mossoroensis* (CMM 4857), *M. nordestinus* (CMM 4846) and *M. semiaridus* (CMM 4830).

### Environmental conditions of the research

The research was conducted under greenhouse conditions in UFERSA, in the period from August 5 to September 23 of 2019, in the municipality of Mossoró RN, Brazil (5°11’15” S, 37°20’39” W). Three trials were carried out and the last results are here presented in this work. The Mossoró climate is very dry and hot with a rainy season that extends from the summer to the fall (Köppen’s classification - BSwh’), with an average temperature of 27.5 °C, average annual rainfall of 670 mm, average air relative humidity of 68.9 %. The temperature ranged from 19 to 34.4 °C during the period of the research.

The trials were conducted in a completely randomized design with five replications, in a 4 × 6 factorial scheme (four melon accessions and six *Monosporascus* species), totaling 24 treatments (pathosystems). The seeds of the accessions were sowed in plastic trays fulfilled with a 2:1 (v/v, sand:Topstrato®) substrate that was autoclaved twice with a 24 h interval at 121 °C and 1.013 × 10^5^ N m^-2^. The same kind of substrate was used to fulfill the 0.5 L pots in which the seedlings were planted.

### Artificial inoculation

The fungal inoculation was made based on the methodology suggested by Pivonia et al. (1997). The inoculum preparation was made with the mycelium of the *Monosporascus* species multiplied in sterilized PDA (potato-dextrose-agar) media. The mycelium grown in PDA media was mixed with a blender in 0.3 L of distilled and sterilized water. To inoculate the plants, 10 mL of each *Monosporascus* species mycelial suspension was pipetted into the substrate of the pots seven days before the seedling transplantation for incubation. The seedlings transplantation was made after 15 days old. The plants were daily irrigated.

### Disease evaluation

The severity evaluation was made 50 days after the transplantation. The roots were removed from the substrate and washed with tap water. The variables analyzed were severity of the disease (rank), with a scores proposed by (Armengol et al. 1998; 1999), and the root dry matter reduction index (RI_DM_).

The rating scores ranged from 0 to 4, with 0: symptomless; 1: mild discoloration (or rot <10 % of the roots); 2: moderate discoloration (or rot of 25 to 35 % of the roots); 3: death of secondary roots (or rot in 50 % of the roots); 4: total necrosis of the roots (or death of the plant). The average reaction was calculated by adding the scores of each accession and divided by the total number of plants evaluated. Consequently, the following classes of genotypes were separated. 0: like immune; 0.1-1.0: highly resistant; 1.1-2.0: moderately resistant; 2.1-3.0: susceptible; 3.1-4.0: highly susceptible (Sales Júnior et al., 2019). The RI_DM_ was calculated by the following expression: RI_DM_ = (DMc - DMi) / (DMc), where DMc: dry matter of access without inoculation (control) and DMi: matter access inoculated with one of the species.

### Data analysis

As the severity variable does not have the residues normally distributed, the original values were transformed according to the Aligned Rank Transform (ART) methodology for non-parametric factor analyses (Wobbrock et al. 2011). A deviance analysis (type III - Wald) was performed for the ranks of the severity variable and analysis of variance (F of Snedecor) for the RI_DM_. To group averages of accessions and the *Monosporascus* species, the Scott-Knott test was used (1 and 5 % probability). Spearman’s correlation coefficient was estimated for verifying the association between severity and the RI_DM_.

The GGE Biplot model was used to evaluate the accession-species interaction (Yan and Kang 2003). The GGE Biplot model holds only the effect of the accessions and the accession-species interaction, and does not separate the effect of the accessions from the accession-species interaction (maintains the multiplicative effect of the factors) and is described as follows: *Y*_*ij*_ −*µ* − *α*_*i j*_− [*β*_*j*_ = *g*_*i*1_*e*_*i*l_ + *g*_*i*2_*e*_*i*2_ + *e*_*ij*_. Where Y_*ij*_ is the performance of the accession *i* on the species *j*; μ is the general average of the observations; *i* is the main effect of the accession *i*; *j* is the main effect of the species *j*; *g*_*i*1_ and *e*_*i*1_ are called the main scores of the accession *i* and the species *j*, respectively; *g*_*i*2_ and *e*_*i*2_ are the secondary scores of the accession *i* and species *j*, respectively; *e*_*ij*_ is the residual error not explained by both effects (‘accession’ or ‘accession-species interaction’).

All analyses were processed with the software R (R Core Team 2020). The approach described by Scott-Knott (1974) was used to group the accessions reaction classes.

## Results

Significant effect of accessions (p <0.01) was observed, indicating heterogeneity between the evaluated plant materials about the severity. But, there was no significant effect of species. There was a significant effect of the accession-species interaction (p <0.01), indicating different behavior of the accessions according to the six *Monosporascus* species inoculated. When interaction between two factors exists, there is no sense on discussing the main factors separately, only their interaction.

The accessions were grouped in two groups when inoculated with the species *M. brasiliensis* (Table 1). The accessions A-16 and ‘HBJ’ made up the first group with the highest average ranks, while C-32 and ‘Goldex’, the second group. Two groups of accessions were made up when inoculated with *M. caatinguensis* and *M. nordestinus*, therein the accession A-16 has the lowest average rank (severity). In addition, the accession A-16 was the most resistant to the *Monosporascus* species evaluated. The accessions made up a single group when inoculated with the species *M. cannonballus, M. mossoroensis* and *M. semiaridus*, respectively.

**Table 1.** Scott-Knott’s test to average ranks and e original averages of the severity of melon accessions inoculated with *Monosporascus* spp. Mossoró RN, 2020.

| Species | Accession |  |  |  | Average |
| --- | --- | --- | --- | --- | --- |
|  | A-16 | C-32 | 'Goldex' | 'HBJ' |  |
| *M. brasiliensis* | 80.5aA | 38.3bA | 53.8bA | 84.8aA | 64.4 |
| *M. cannonballus* | 29.5aB | 55.9aA | 65.0aA | 47.1aB | 49.4 |
| *M. caatinguensis* | 14.5bB | 78.1aA | 78.1aA | 59.8aB | 57.7 |
| *M. mossoroensis* | 50.9aA | 71.4aA | 64.7aA | 48.8aB | 61.2 |
| *M. nordestinus* | 14.5bB | 62.6aA | 80.2aA | 96.0aA | 63.3 |
| *M. semiaridus* | 71.4aA | 55.9aA | 78.1aA | 70.0aB | 68.9 |
| Average | 45.1 | 60.4 | 69.9 | 67.9 |  |
| Effect | df<sub>1</sub> | df<sub>2</sub> | Test (Type III – Wald) |  |  |
| | | | F | $\chi^2$ | |
| Accessions (A) | 3 | 96 | 4.84<sup>**</sup> | 14.53<sup>**</sup> |  |
| Species (S) | 5 | 96 | 1.08<sup>ns</sup> | 5.42<sup>ns</sup> |  |
| A x S | 15 | 96 | 3.01<sup>**</sup> | 45.07<sup>**</sup> |  |

<sup>\*\*</sup>, <sup>\*</sup>: Significant at (p<0,01) and (p<0,05) by the tests F and Chi-square ( $\chi^2$ ); df<sub>1</sub>: degrees of freedom of the numerator; df<sub>2</sub>: degrees of freedom of the denominator. 'HBJ': Hales Best Jumbo. Averages followed by the same lowercase letters in the row belongs to the same group according to the Scott-Knott's approach (p>0.05).

The species formed two groups when they were inoculated in the accession A-16 (Table 1). The species *M. brasiliensis, M. mossoroensis* and *M. semiaridus* made up the first group and were more aggressive (higher average rank). When inoculated in ‘HBJ’, the species were set apart in a first group composed of *M. brasiliensis* and *M. nordestinus*. These two species were more aggressive to that accession (‘HBJ’). Then, the other species were added in the second group. When inoculated at the C-32 and ‘Goldex’ accession, the species formed a single group, respectively.

The GGE Biplot methodology can be used to approach the non-additive effect (interaction) between pathogens and plant hosts (Yan and Kang 2003). In the Biplot graph, the two axes together explained 97.25 % of the total variation in the accession-species interaction. The polygon formed was composed of four vertices (Figure 1A). In each vertex there is an inoculated accession, which has the highest average rank in its respective sector, being, therefore, the one with the most severity.

**Figure 1.**
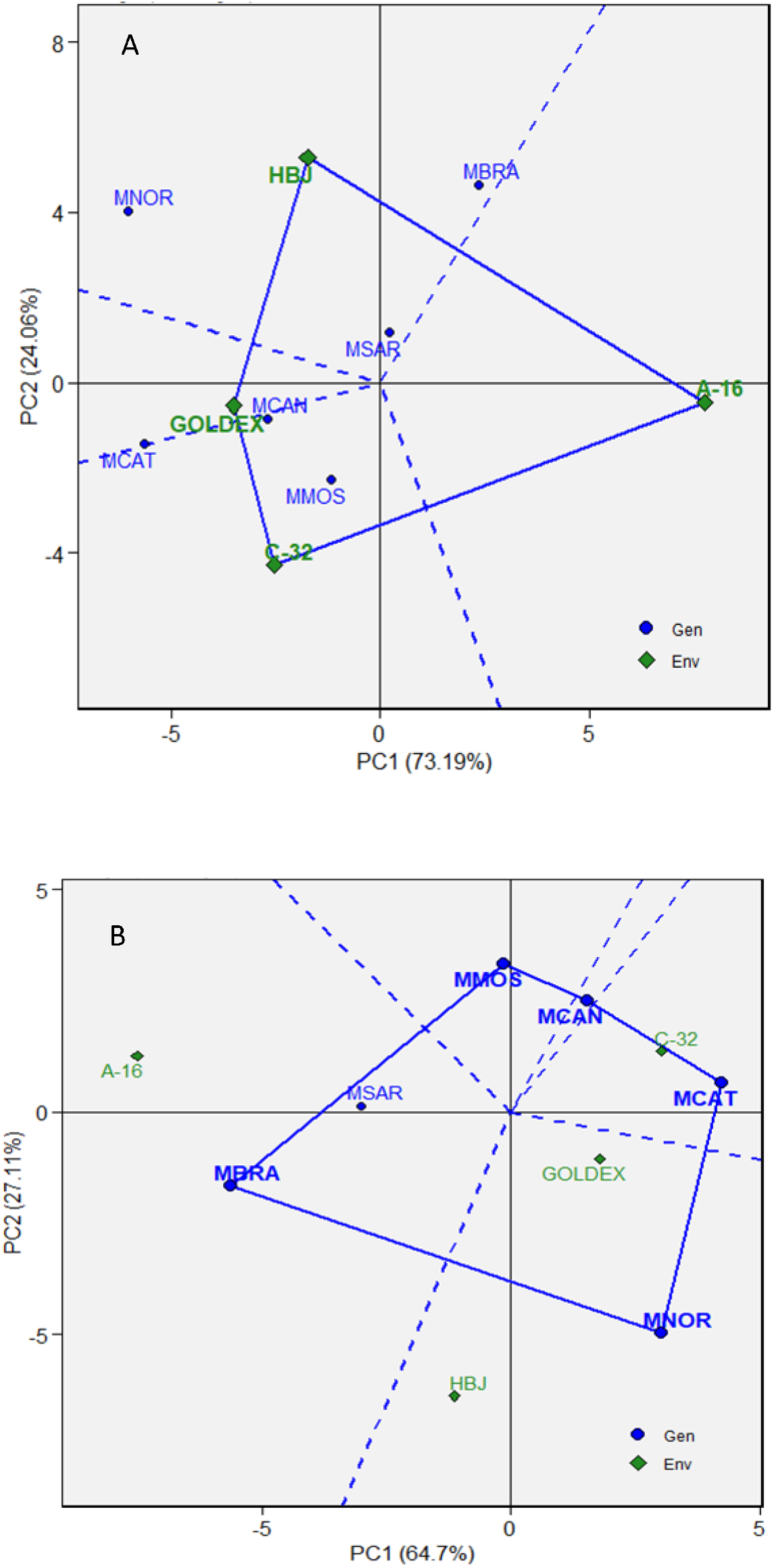
(A) GGE Biplot with the distribution of the genotypes (on the vertices) inoculated with six species of the genus *Monosporascus*. (B) GGE Biplot with the distribution of the species of the genus *Monosporascus* (on the vertices) inoculated in four accession of melon. MBRA: *M. brasiliensis*, MCAN: *M. cannonballus*, MCAT: *M. caatinguensis*, MMOS: *M. mossoroensis*, MNOR: *M. nordestinus* e MSAR: *M. semiaridus*.

Counterclockwise, in the first vertex was the accession ‘HBJ’, that the species *M. nordestinus, M. brasiliensis* and *M. semiaridus* were more virulent in relation to the others. At the second vertex was the accession ‘Goldex’, which has higher average rank in relation to *M. caatinguensis*. The accession C-32, at the third vertex, was associated with the highest average rank (highest severity) when inoculated with *M. cannonballus* and *M. mossoroensis*. The accession A-16, at the fourth vertex, did not interact with the species and showed less severity (higher performance).

The GGE Biplot modeling also can be set in the opposite direction, i.e., setting the species in the vertices (Figure 1B). The six species were set into five vertices of the polygon of the GGE Biplot graph. In the counterclockwise direction, the species *M. brasiliensis*, in the second vertex, interacted more with the accession *A-16*, while *M. nordestinus*, in the third vertex, interacted with ‘Goldex’ and ‘HBJ’. *M. caatinguensis*, located in the fourth vertex, interacted only with the access C-32.

The species *M. caatinguensis* performed better (greater virulence) because it was more to the right-hand side on the x-axis, i.e., it was the most adapted species (due to the proximity in the graph) to the accession C-32. On the contrary, the species *M. brasiliensis* performed worst (less virulence) because it was more to the left-hand side on the x-axis, followed by *M. semiaridus*.

The accessions were classified according to the severity of reaction of the six inoculated species (Figure 2). The accession A-16 was highly resistant to *M. nordestinus, M. caatinguensis* and *M. cannonballus*, moderately resistant to *M. mossoroensis* and *M. semiaridus*, but susceptible to *M. brasiliensis*. The accession C-32 was susceptible to the species *M. caatinguensis*, but moderately resistant to the other species. The accession ‘Goldex’ was susceptible to the species *M. caatinguensis, M. nordestinus* and *M. semiaridus*, and moderately resistant to *M. brasiliensis, M. cannonballus* and *M. mossoroensis*. The accession ‘Hales Best Jumbo’ (‘HBJ’) was susceptible to *M. brasiliensis* and *M. nordestinus* and moderately resistant to the other species.

**Figure 2.**
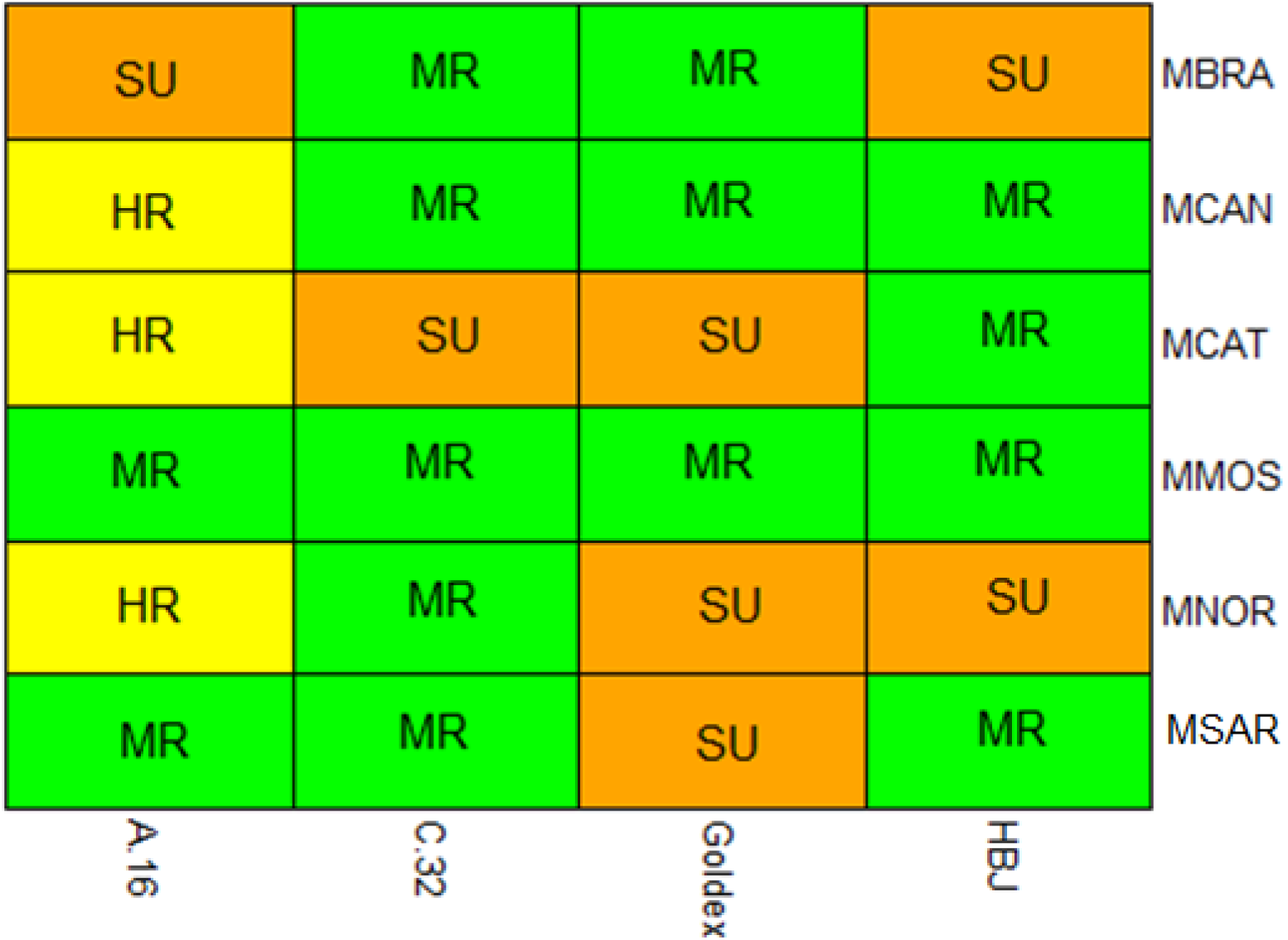
Melon accessions reaction to six species of the genus *Monosporascus*. HR: highly resistant, MR: moderately resistant, SU: susceptible. ‘HBJ’: Hales Best Jumbo. MBRA: *M. brasiliensis*, MCAN: *M. cannonballus*, MCAT: *M. caatinguensis*, MMOS: *M. mossoroensis*, MNOR: *M. nordestinus* e MSAR: *M. semiaridus*.

The accessions and species did not present effect for the RI_DM_, but for the accession-species interaction (Table 2). When inoculated with the species *M. brasiliensis*, the accessions were grouped into two groups, the first group composed only by A-16 and the second by the other accessions. The accessions were divided into two groups when inoculated by the species *M. cannonballus*. The first group formed by the accessions of the greatest RI_DM_, the accession ‘Goldex’, and the second group formed by the other accessions. When inoculated by the species *M. mossoroensis*, the accession with the smallest RI_DM_ was A-16 while the other accessions with the highest average, were gathered into a single group. The accessions were gathered in two groups when inoculated by the species *M. nordestinus*. The group with the highest averages was composed of the accessions ‘Goldex’ and ‘HBJ’. The accessions were allocated to the same group when inoculated by the species *M. caatinguensis* and *M. semiaridus*.

**Table 2.** Unfolding of the root matter reduction index (RI_DM_) of melon accession inoculated with *Monosporascus* spp. Mossoró RN, 2020.

| Species | Accession |  |  |  |
| --- | --- | --- | --- | --- |
|  | A-16 | C-32 | ‘Goldex’ | ‘HBJ’ |
| | $RI_{DM}$ (g) | | | |
| *M. brasiliensis* | 0.53aA | 0.38bA | 0.45bA | 0.26bB |
| *M. cannonballus* | 0.30bB | 0.24bA | 0.58aA | 0.10bB |
| *M. caatinguensis* | 0.17aB | 0.17bA | 0.32aB | 0.27aB |
| *M. mossoroensis* | 0.24bB | 0.46aA | 0.40aA | 0.36aB |
| *M. nordestinus* | 0.26bB | 0.21bA | 0.46aA | 0.59aA |
| *M. semiaridus* | 0.39aB | 0.29aA | 0.25aB | 0.23aB |

| Effect | df<sub>1</sub> | df<sub>2</sub> | Test (F of Snedecor) |
| --- | --- | --- | --- |
| Acessions (A) | 3 | 96 | 2.01<sup>ns</sup> |
| Species (S) | 5 | 96 | 2.25<sup>ns</sup> |
| A x S | 15 | 96 | 1.91<sup>*</sup> |

\*\*, \*: Significant to ( $p < 0.01$ ) e ( $p < 0.05$ ) bu the test of Snedecor; df<sub>1</sub>: degrees of freedom of the numerator; df<sub>2</sub>: degrees of freedom of the denominator. ‘HBJ’: Hales Best Jumbo. Averages followed by the same lowercase letters in the row belongs to the same group according to the Scott-Knott’s approach ( $p > 0.05$ ).

About the accession A-16, the species *M. brasiliensis* was the most virulent causing a reduction of more than 50 % in the A-16 root dry matter, being allocated in a separate group from the other species. The species were gathered in the same group when inoculated the accession C-32. The species *M. cannonballus, M. brasiliensis, M. mossoroensis* and *M. nordestinus*, gathered in the same group, were more virulent about the accession ‘Goldex’. The species were combined into two groups when inoculated over the accession ‘HBJ’. The first group formed by the species *M. nordestinus*, more virulent, while the second group formed by the other species.

Spearman’s correlation coefficients were estimated between severity and RI_DM_ within each accession and each species. Significant values were found in practically all accessions and species, but there were some exceptions: the accession C-32 and the species *M. brasiliensis* and *M. semiaridus* (Figure 3). The positive sign indicates the two variables grow in the same direction (directly proportional). When the severity increases there is a reduction in the dry matter of the plant. The correlation, considering all the data, also followed the same fashion, positive and significant value (r = 0.58 *).

**Figure 3.**
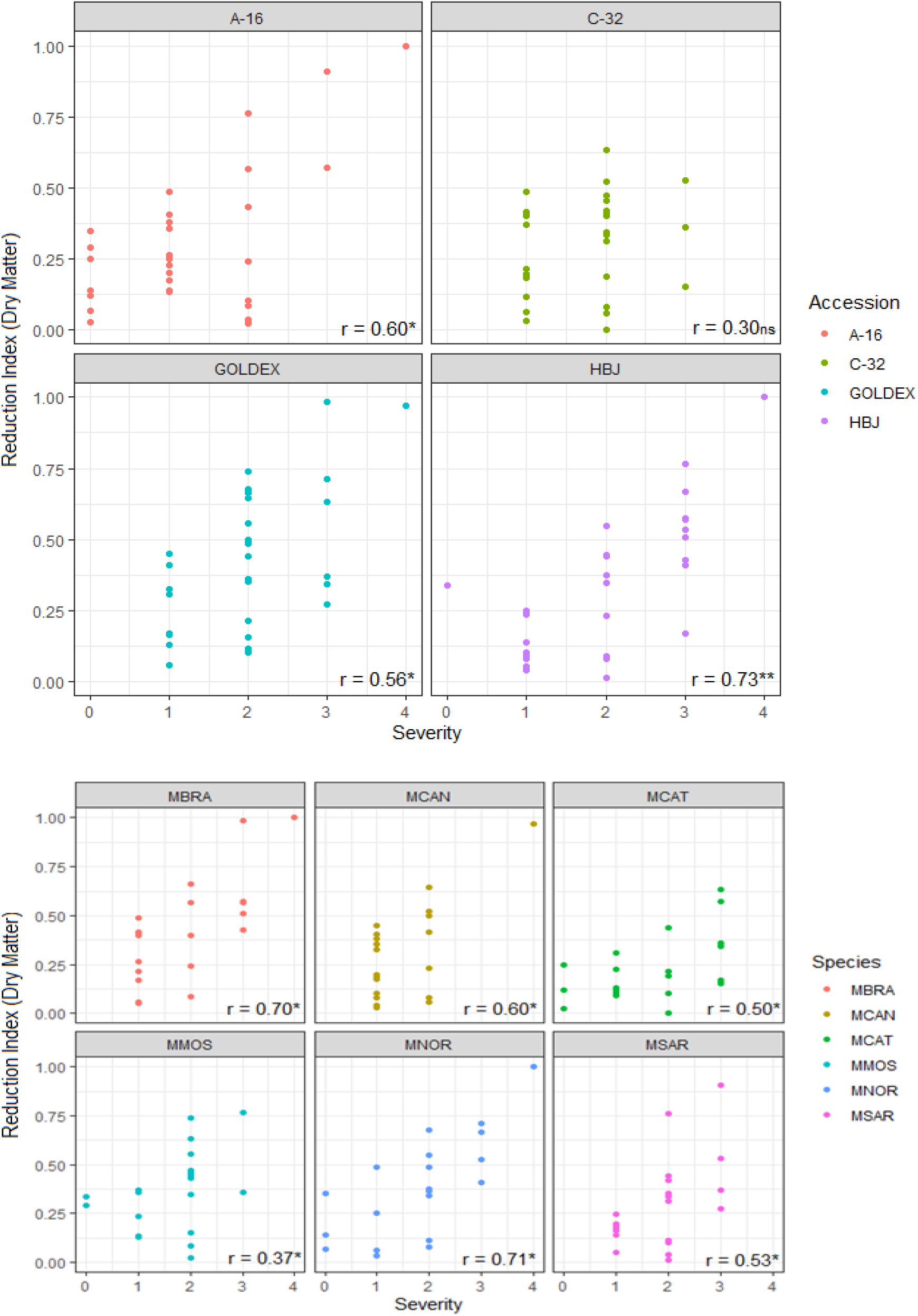
Estimations of the coefficient of Spearman’s correlation between the severity and the root matter reduction index (RI_DM_) among each accession and species. Mossoró RN, 2020. MR: moderately resistant, SU: susceptible. ‘HBJ’: Hales Best Jumbo. MBRA: *M. brasiliensis*, MCAN: *M. cannonballus*, MCAT: *M. caatinguensis*, MMOS: *M. mossoroensis*, MNOR: *M. nordestinus* e MSAR: *M. semiaridus*.

## Discussion

To date, there is no reports on the virulence of other species of *Monosporascus* in melon genotypes. It was only known that MRRVD in melon and watermelon is caused by the species *M. cannonballus* and *M. eutypoides* worldwide (Castro et al. 2020). Despite this, five novel species of *Monosporascus* isolated from weeds root *Boerhavia diffusa* L. and *Trianthema portulacastrum* Linn., usually found in melon production fields in northeastern Brazil (Negreiros et al. 2019) were identified. The present work is the first report of the pathogenicity and virulence of the species *M. brasiliensis, M. caatinguensis, M. mossoroensis, M. nordestinus* and *M. semiaridus* in melon. This data is very important because these species are potentially disease-causing for melon.

Another important look of this work is that the species have a different behavior due to accessions. Despite this, it was found that the species *M. cannonballus*, notoriously the agent of MRRVD worldwide (Sarpeleh 2008; Al-Mawaali et al. 2013; Yan et al. 2016; Markakis et al. 2018; Sales Júnior et al. 2019) and the novel species *M. mossoroensis* was the least virulent since none of the accessions was classified as susceptible to both species. Particularly, the accession A-16 was highly resistant to *M. cannonballus*, and of moderate or high resistance when inoculated with each of these species. On the other hand, the other species of *Monosporascus* caused a susceptibility reaction in at least one of the accessions (Figure 2). It is noteworthy that a reduced number of accessions were used in the study, therefore, more studies including a greater number of genotypes are needed.

Regarding the reaction of the genotypes in this work, it is worth of mentioning that all are resistant or moderately resistant to the species *M. cannonballus*. This species was identified in Brazil in melon fields (Sales Júnior et al. 2003) and later in watermelon fields (Sales Júnior et al. 2010). It is one of the most isolated species in melon roots in the Brazilian semiarid (Marinho et al. 2002) together with *Macrophomina phaseolina* and *Fusarium solani* (Ambrósio et al. 2015). Therefore, the identification of resistant accessions to *M. cannonballus* is relevant. Some sources with moderate resistance such as F35a, P6a (Cohen et al. 1996), ‘Sfidak khatdar’ and ‘Sfidak bekhat’ (Salari et al. 2012), as well as high resistance; 20608, 20747, 20826 (Crosby 2001), were identified.

In Spain, under field conditions and in greenhouse artificial inoculation, the accession Pat 81 (ssp. *agrestis*) was identified with a high level of tolerance (Esteva and Nuez 1994; Iglesias and Nuez 1998, Iglesias et al. 1999; 2000a, b). This plant material was used in a breeding program for backcrosses that generated resistant lines of melon *Piel de Sapo* (Fita et al. 2009a, b). The accession C-32 used in this work is Pat 81, which presented moderate resistance to five species, but was susceptible to *M. caatinguensis*. The results of the present study confirm the importance of this material as a source of resistance, however, it reports, for the first time, its susceptibility to a species of *Monosporascus*.

The accession ‘Hales Best Jumbo’ (‘HBJ’) was moderately resistant to the species *M. cannonballus* as well as, *M. caatinguensis, M. mossoroensis* and *M. semiaridus*. In one of the first efforts to identify sources of resistance to *M. cannonballus*, Mertely (1993) concluded that Hales Best Jumbo, Honey Dew Green Flesh, Improved, Cruiser, Durango, PI 12411 and Laredo were tolerant to this fungal species. On the other hand, ‘HBJ’ was susceptible to the species *M. brasiliensis* and *M. nordestinus*.

The hybrid ‘Goldex’ was susceptible to the species *M. caatinguensis, M. nordestinus* and *M. semiaridus* (Figure 2). Furthermore, we highlight that the accession ‘Goldex’ presented high values of the RI_DM_ (Table 2). This hybrid is also highly susceptible to powdery mildew (*Podosphaera xanthii*) and the leafminer (*Liriomyza sativa*) but is still the preferred one due to its high quality and post-harvest fruit. It is estimated that most of the melon area in the Brazilian semiarid (around ∼ 22,000 ha / year) has been cultivated with ‘Goldex’ in the past 15 years. The work information shows the potential fragility of the hybrid ‘Goldex’. On the other hand, a strategy for future improvement is to obtain yellow melon cultivars resistant to the main pathogens and with a ‘Goldex’ background.

The accession A-16 was the most promising one since it was highly resistant to three species (*M. cannonballus, M. caatinguensis* and *M. nordestinus*) and moderately resistant to two (*M. mossoroensis* and *M. semiaridus*). In addition, it has lower RI_DM_ (Table 2). The accession A-16 has fruits with high mesocarp firmness, high titratable acidity, low content of soluble solids (Dantas et al. 2015) and resistance to *Myrothecium roridum* (Nascimento et al. 2012). This accession belongs to the *acidulus* botanical group, very usual in India, which has provided many sources of resistance to pathogens such as fungi, viruses and insects (Dhillon et al. 2012).

A worthy aspect to be highlighted in breeding programs is that the tolerance in melon to *M. cannonballus* is strictly related to the root system. In the present work, only the RI_DM_ of the genotypes was estimated. There was a positive and significant association between the severity and the RI_DM_, that is, the greater the severity, the lower the dry matter of the roots of the inoculated plants when compared with the non-inoculated plants (Figure 3). Crosby et al. (2000) observed that resistant cultivars have higher averages for the total root length, the average root diameter, the number of root branches, the number of thin roots (0.0-0.5 mm) and the number of small roots (0.5-1.0 mm) compared to susceptible cultivars. The tolerance of the C-32 accession is explained by the high vigor and pronounced branching of its root system. This accession has high root mass, even when infected, when compared to the susceptible cultivar Pioñet (Dias et al. 2002).

It should be noted that the inoculation performed in the present study used a mycelial suspension, since some species of *Monosporascus* could not be induced to sporulation (Negreiros et al. 2019). In fact, several methodologies can be developed to evaluate the virulence of pathogens, by observing the severity of attacks on plant tissues, such as damage to the hypocotyl, in primary and secondary roots, reduction of leaf area (Bruton et al. 2000). In these pathogenicity studies, several inoculation methods can be adopted, such as inoculating the soil with agar colonized with the fungus (Uematsu et al. 1985; Tsay and Tung 1995; Martyn and Miller 1996; Pivonia et al. 1997), oat husks mixed with sand in pots (Mertely 1993; Karlatti et al. 1997; Pivonia et al. 1997), sorghum grains colonized by fungi and incorporated into the soil (Lovic et al. 1994) etc. These studies proved the effectiveness of the different inoculation methods.

The present work represents the first study of pathogenicity of *Monosporascus* species described in Brazil in comparison with *M. cannonballus*, the species of this genus most distributed and studied worldwide.

The accession A-16 was identified as resistant to most species. In addition, the breeding value of the accession C-32 as a source of resistance was confirmed. These accessions can be used in classic breeding programs and, or, as rootstocks. Still with respect to C-32, it was used as a source of resistance in a backcross program that generated resistant or tolerant lines of *Piel de Sapo* to *M. cannonballus* and with excellent quality fruits (> 12 % of total soluble solids) (FITA et al. 2009a, b). In the second alternative, C-32 was used as a rootstock with excellent results. Plants grafted onto the accession C-32 had few symptoms and good fruit quality when compared to plants grafted onto rootstocks of the genus *Cucurbita*. In addition, no incompatibility was observed when using the accession C-32 as a rootstock (Fita et al. 2007).

## Declarations

## Acknowledgements

Coordination of Higher Education Personnel Improvement – CAPES, for providing the master degree scholarship for the first author.

